# Population position along the fast-slow life-history continuum predicts intraspecific variation in actuarial senescence

**DOI:** 10.1101/621425

**Authors:** Hugo Cayuela, Jean-François Lemaître, Eric Bonnaire, Julian Pichenot, Benedikt R. Schmidt

**Affiliations:** Département de Biologie, Institut de Biologie Intégrative et des Systèmes (IBIS), Université Laval, Pavillon Charles-Eugène-Marchand, Québec, QC G1V 0A6, Canada; Université Lyon 1, CNRS, UMR 5558, Laboratoire de Biométrie et Biologie Evolutive, F-69622, Villeurbanne, France; Office National des Forêts, Agence de Verdun, route de Metz, 55100, Verdun, France; URCA, CERFE, Centre de Recherche et Formation en Eco-éthologie, 08240, Boult-aux-Bois, France; Institut für Evolutionsbiologie und Umweltwissenschaften, Universität Zürich, Winterthurerstrasse 190, 8057 Zürich, Switzerland; Info Fauna Karch, UniMail, Bâtiment G, Bellevaux 51, 2000 Neuchâtel, Switzerland

**Keywords:** senescence, life history, fast-slow continuum, amphibians, *Bombina variegata*

## Abstract

Patterns of actuarial senescence can be highly variable among species. Previous comparative analyses revealed that both age at the onset of senescence and rates of senescence are linked to the species’ position along the fast-slow life-history continuum. As there are few long-term datasets of wild populations with known-age individuals, intraspecific (i.e. between-population) variation in senescence is understudied and limited to comparisons of wild and captive populations of the same species, mostly birds and mammals. In this paper, we examined how population position along the fast-slow life history continuum affects senescence patterns in an amphibian, *Bombina variegata*. We used capture-recapture data collected in four populations with contrasted life history strategies. Senescence trajectories were drawn using Bayesian capture-recapture models. We show that in “slow” populations the onset of senescence was earlier and individuals aged at a faster rate than individuals in “fast” populations. Our study provides one of the few empirical examples of between-population variation in senescence patterns in the wild and confirms that the fast-slow life history gradient is associated with both macroevolutionary and microevolutionary patterns of senescence.

## Introduction

Most commonly accepted evolutionary theories of ageing posit that survival should decline with increasing age in any age-structured population (Hamilton 1966), a demographic process coined as actuarial senescence. Medawar’s (1952) mutation accumulation theory first stated that organisms age because the strength of natural selection weakens with age after first reproduction and therefore there is no purging of deleterious mutations that are only expressed late in life. In addition, actuarial senescence can emerge as a by-product of natural selection through antagonistic pleiotropy (Williams 1957). An allele may confer a benefit to the bearer early in life but may also be responsible for an impaired survival later in life. Finally, the disposable soma theory of aging postulates that actuarial senescence can result from a trade-off between allocation to reproduction in early life and somatic maintenance (Kirkwood 1977, Kirkwood & Austad 2000). In other words, individuals that preferentially allocate resources to growth and/or reproduction (e.g., gamete production and parental care) will have much less resources for somatic maintenance (e.g., enzyme-based repair mechanisms), which will ultimately lead to a decline in performance of fitness-related traits (e.g., survival) at advanced ages. So far, most studies focused on aging in the wild have been embedded within the two theoretical frameworks offered by both antagonistic pleiotropy and disposable soma theories of aging as these two theories share the similar prediction of a trade-off between reproductive effort and senescence (Gaillard & Lemaître 2017).

Although a few case studies on free-ranging populations failed to detect any increase in mortality rate with age (Jones et al. 2014), most recent syntheses revealed two major facts. First, actuarial senescence is a nearly ubiquitous process in the living world (Nussey et al. 2013, Shefferson et al. 2017). Second, patterns of senescence can be highly variable among species (Jones et al. 2014, Tidière et al. 2016, Colchero et al. 2019). Comparative analyses have shown that senescence patterns across multi-cellular organisms can be predicted by ecological traits, lifestyles and covariation among life-history traits (Péron et al. 2010, Ricklefs 2010, Gaillard et al. 2016, Salguero-Gómez & Jones 2017). In particular, both age at the onset of senescence and rates of senescence appear to be linked to the position of a species along the fast-slow life-history continuum. Organisms that occupy the fast end of the continuum – short generation time, high annual fecundity, and low mean survival rates (Stearns 1983) – tend to experience earlier and faster senescence than organisms at the slow end of the continuum (Jones et al. 2008a, Salguero-Gómez & Jones 2017). Since life history variation does not only occur among, but also within species (Berven & Gill 1983, Cayuela et al. 2017), intraspecific variation in senescence patterns is expected and can be selected for (Stearns 2000, Stearns et al. 2000, Brommer et al. 2007, Holand et al. 2017). In particular, we expect intraspecific variation along a fast-slow life history continuum whenever mortality rates differ among populations (Stearns 2000, Stearns et al. 2000). Within the same species, factors such as climate (Nevoux et al. 2010), habitat predictability (Cayuela et al. 2016a), predation (Nilsen et al. 2009) and pathogens (Jones et al. 2008b) may affect growth, fecundity and age-specific survival in ways that affect the position of a given population along the fast-slow continuum. Insights into the intraspecific (i.e. between-population) variability in actuarial senescence patterns have been obtained almost exclusively through the comparisons of captive and wild populations of birds and mammals or through selection experiments using *Drosophila* (Stearns et al. 2000, Ricklefs & Scheuerlein 2001, Lemaître et al. 2013, Tidière et al. 2016; but see Austad 1993 and Holand et al. 2016 for case studies in Virginia opossums, *Didelphis virginiana* and house sparrows, *Passer domesticus*, respectively), which is notably due to the scarcity of long-term individual data associated with age records from multiple free-ranging populations.

Most of the studies on senescence in vertebrates focus on endothermic organisms, such as birds and mammals, and largely neglect ectotherms (Bronikowski et al. 2002, Jones et al. 2014, Lemaître et al. 2015, but see Sparkman et al. 2007, Massot et al. 2011, Warner et al. 2016). Both actuarial and reproductive senescence are understudied in ectotherms, mainly due to the scarcity of demographic data and methodological limitations (juveniles cannot be marked easily and information about the age of individuals is often partial and uncertain). However, ectotherms such as amphibians could be also excellent biological models for ageing studies, especially to investigate the evolution of senescence, as they display broad variation of life history traits (Turner 1962, Werner 1986, Van Bocxlaer et al. 2010): while some species are iteroparous with lifespans exceeding ten years, others live only one or two years and are essentially semelparous (Wells 2010). Moreover, intraspecific variation of demographic parameters exists along environmental gradients (Berven & Gill 1983, Morrison & Hero 2003, Sinsch et al. 2010), which has resulted in a fast-slow life history gradient (Cayuela et al. 2017).

We examined how variation in population position along the fast-slow continuum affects actuarial senescence patterns in an amphibian, the yellow-bellied toad (*Bombina variegata*). In Western Europe, the species displays two life history strategies associated with contrasted habitats (Cayuela et al. 2016a, 2016c). In river environments, individuals have slower life histories than in forest environments: they have higher juvenile and adult survival rates, more canalized survival rates with a lower temporal variance and a lower annual recruitment (i.e., number of juveniles produced per female per year) (Cayuela et al. 2016a; Fig. 1). Here, we analysed age-specific mortality rates using capture-recapture data and Bayesian survival trajectory analyses (Colchero et al. 2012). We specifically investigated how actuarial senescence patterns (i.e., the onset of senescence and the speed and the shape of the relationship between mortality rate and age) differed in populations with slow and fast life histories. We focused our analyses on post-metamorphic survival as we did not expect senescence during larval development and assumed that senescence can begin once structural development is completed (i.e., metamorphosis). We expected that the pattern of intraspecific variation in senescence among populations would be similar to pattern in variation among species. Therefore, we tested the prediction that a fast pace-of-life should be associated with an earlier, faster senescence among populations of *B. variegata*.

**Fig. 1.**
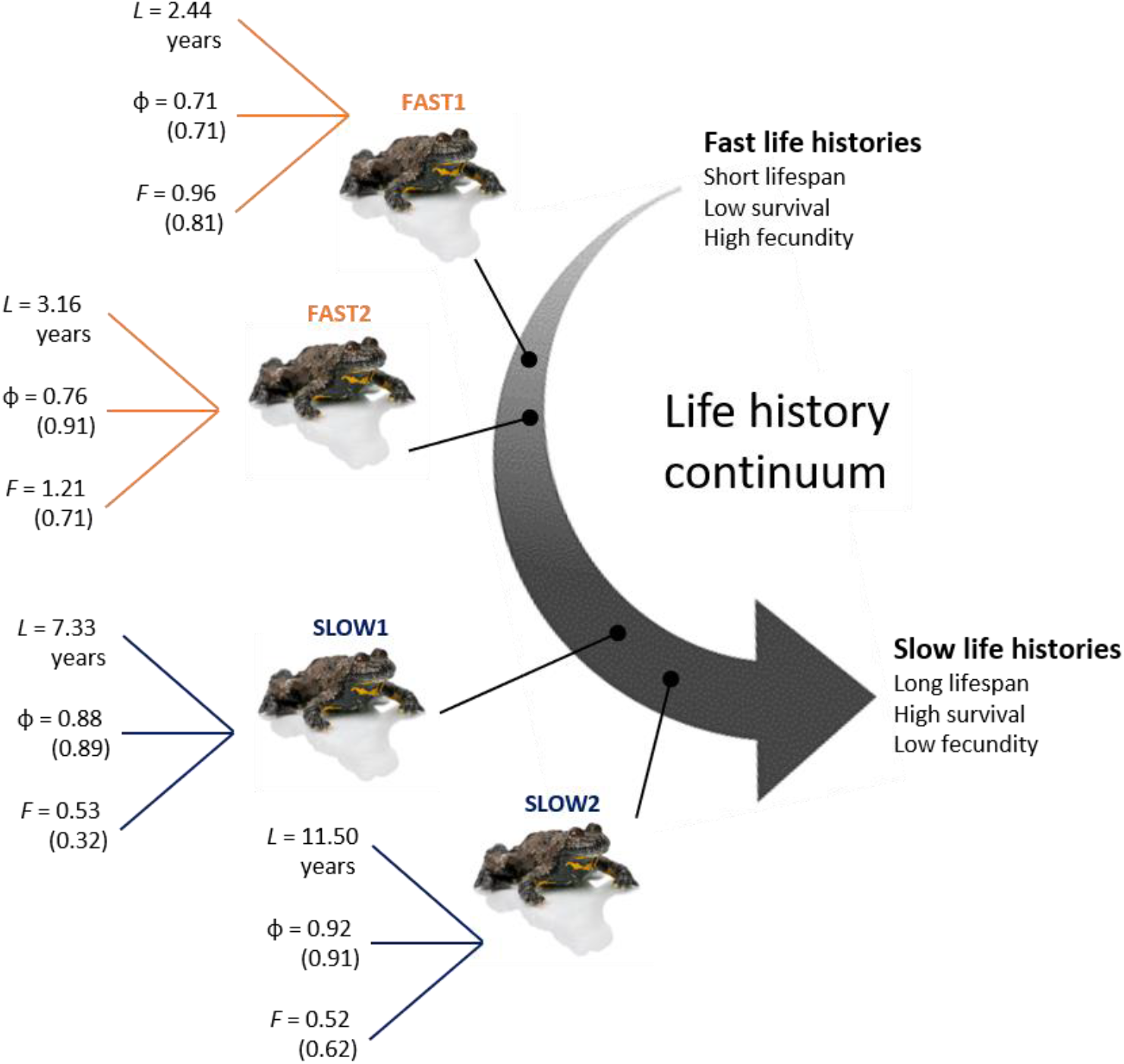
Intraspecific variation along the slow-fast life-history gradient in *Bombina variegata* populations: life expectancy (L), annual adult survival probability (φ), recruitment (F) in terms of number of juveniles produced by female per year. Coefficient of variation are provided in brackets. The data provided in the figure are extracted from Cayuela et al. (2016a, 2016b).

## Material and methods

### Study system

*Bombina variegata* populations occur in different types of habitats where the spatiotemporal pattern of breeding resource availability differs widely. In river environments, the patches of rock pools used by toads to reproduce are constantly available in space and time, making breeding resources highly predictable at the scale of the lifetime of an individual. By contrast, rut patches resulting from logging operations appear and disappear stochastically in forests, making breeding resource availability unpredictable. The annual probability of patch appearance varies from 0.20 to 0.50 while the rate disappearance ranges from 0.05 to 0.20, depending on the year and the population (Cayuela et al. 2016a).

Here, we used mark-recapture data (Appendix 1) from two populations with fast life history strategies (SLOW1 and SLOW2) and two populations with slow life history strategies (FAST1 and FAST2) in France (Appendix 1, Fig. 1). In each population, individuals were surveyed during the breeding season, from 5 – 9 years using capture-recapture methodology (Cayuela et al. 2016a, 2016c). The number of identified individuals was 1154 at SLOW1, 768 at SLOW2, 580 at FAST1 and 9418 at FAST2. The number of captures was 2907 at SLOW1, 1980 at SLOW2, 949 at FAST1 and 12780 at FAST2. In this species, individuals cannot be surveyed before an age of one because the ventral color pattern used to identify individuals is not fixed before that age (Cayuela et al. 2016a, 2016c). Moreover, age can be determined only during the 2 years following metamorphosis (Cayuela et al. 2016b). Sex cannot be assessed with certainty before sexual maturity (2-3 years old) due to the lack of nuptial pads in immature males. For this reason, sex was not considered in our analyses. Yet, we expect that excluding the sex from our analyses should not alter our conclusions since previous studies in these populations did not detect sex-specific effect on survival (Cayuela et al. 2016a, 2016c). Details about the number of survey years, individual age and sampling effort are provided in Appendix 1 (Table 1). A more complete description about the capture-recapture survey, the individual recognition method and population description can be found in Cayuela et al. (2016a, 2016c).

**Table 1.** Deviance information criterion (DIC) for each of the mortality function considered in the four studied populations of Bombina variegata. We considered the four mortality functions implemented in BaSTA program: exponential (EXP), Gompertz (GOM), Weibull (WEI) and logistic (LOG). For the three last functions, we considered three potential shapes: simple that only uses the basic functions described above (“simple”); Makeham (“make”); and bathtub (“bath”).

| Function | SLOW1 | SLOW2 | FAST1 | FAST2 |
| --- | --- | --- | --- | --- |
| LOG_simple | 13883.1 | 9770.011 | 6539.856 | 79770.16 |
| LOG_bath | 13408.63 | 9511.085 | **6164.568** | **76039.06** |
| LOG_make | 13878.78 | 9706.06 | 6559.085 | 79669.19 |
| EXP | 16151.69 | 11085.02 | 8179.035 | 120984.4 |
| GOM_simple | 13515.9 | 9104.43 | 6859.955 | 91054.45 |
| GOM_bath | **13210.67** | **8855.467** | 6797.639 | 89020.97 |
| GOM_make | 13534.36 | 9065.034 | 6888.318 | 91053.47 |
| WEI_simple | 13793.82 | 9451.142 | 6811.184 | 87726.47 |
| WEI_bath | 13406.18 | 9230.966 | 6653.47 | 85057.55 |
| WEI_make | 13790.13 | 9464.459 | 6805.522 | 87757.15 |

### Capture-Recapture modeling

We investigated actuarial senescence patterns using Bayesian survival trajectory analyses implemented in the R package BaSTA (Colchero et al. 2012a, 2012b). BaSTA allowed us to account for imperfect detection, left-truncated (i.e., unknown birth date) and right-censored (i.e., unknown death date) capture-recapture data in our analysis. Our analyses focus on the post-metamorphic stage at which senescence is expected to occur (as in Colchero et al. 2019). It allows estimation of two parameters: age-dependent survival and the proportion of individuals dying at a given age (i.e. age-dependent mortality rate). Given the results of previous analyses (Cayuela et al. 2016a, 2016b), we allowed recapture probabilities to vary among years. As the study period and number of survey years differ among populations (Appendix 1), the four populations were analyzed separately. We used DIC to select models that fitted the data best (Colchero et al. 2019) and we compared the outputs of the best-supported model of the four populations by inspecting mean estimates and 95% CI (Anderson et al. 2001, Amrhein et al. 2019). This allowed us to investigate population-specific variation in the shape of the age-specific mortality patterns. We considered the four mortality functions implemented in BaSTA: exponential, Gompertz, Weibull and logistic. For the three last functions, we considered three potential shapes: *simple* that only uses the basic functions described above; *Makeham* (Pletcher 1999); and *bathtub* (Silver 1979). As individuals cannot be individually recognized before one year old in *B. variegata*, we conditioned the analyses at a minimum age of one. Four MCMC chains were run with 50000 iterations and a burn-in of 5000. Chains were thinned by a factor of 50. Model convergence was evaluated using the diagnostic analyses implemented in BaSTA, which calculate the potential scale reduction for each parameter to assess convergence. For all populations, we used DIC to compare the predictive power of each mortality function and its refinements (Spiegelhalter et al. 2002, Colchero et al. 2013, 2019). Simulations by Colchero & Clark (2012) showed that BaSTA models are highly efficient even when dates of birth and death are unknown and the study is short (i.e. the study length is less than the mean life expectancy). They also showed that estimates are more accurate when recapture rate is high. Therefore, we are confident that our inference is robust as recapture rates are relatively high (between 0.54 in FAST1 to 0.84 in SLOW2), birth time uncertainty is moderate (between 52% and 57% of known-age individuals in our sampled populations).

## Results

Our analyses revealed contrasting patterns of age-specific survival and mortality rate between individuals from populations with slow and fast life histories. The shape of the relationship between mortality rate and age differed broadly between the two life history strategies. Based on DIC comparisons (Appendix 1, Table 2), the relationship was best described by a Gompertz function with a bathtub shape in the “slow” populations (Fig. 2A and 2B) while a logistic function with a bathtub shape provided the best fit in both “fast” populations (Fig. 2C and 2D). In “slow” populations, mortality rate slowly increased with age until 8-10 years after which it increased quickly (for model parameter estimates, see Appendix 1, Table 3–4). This indicates a late, slow senescence and a long reproductive lifespan. By contrast, in “fast” populations, mortality rate increased rapidly after individuals are 2-3 years old and reached an asymptote at 3 to 4 years. This suggests an early, fast senescence with a short reproductive lifespan.

**Fig. 2.**
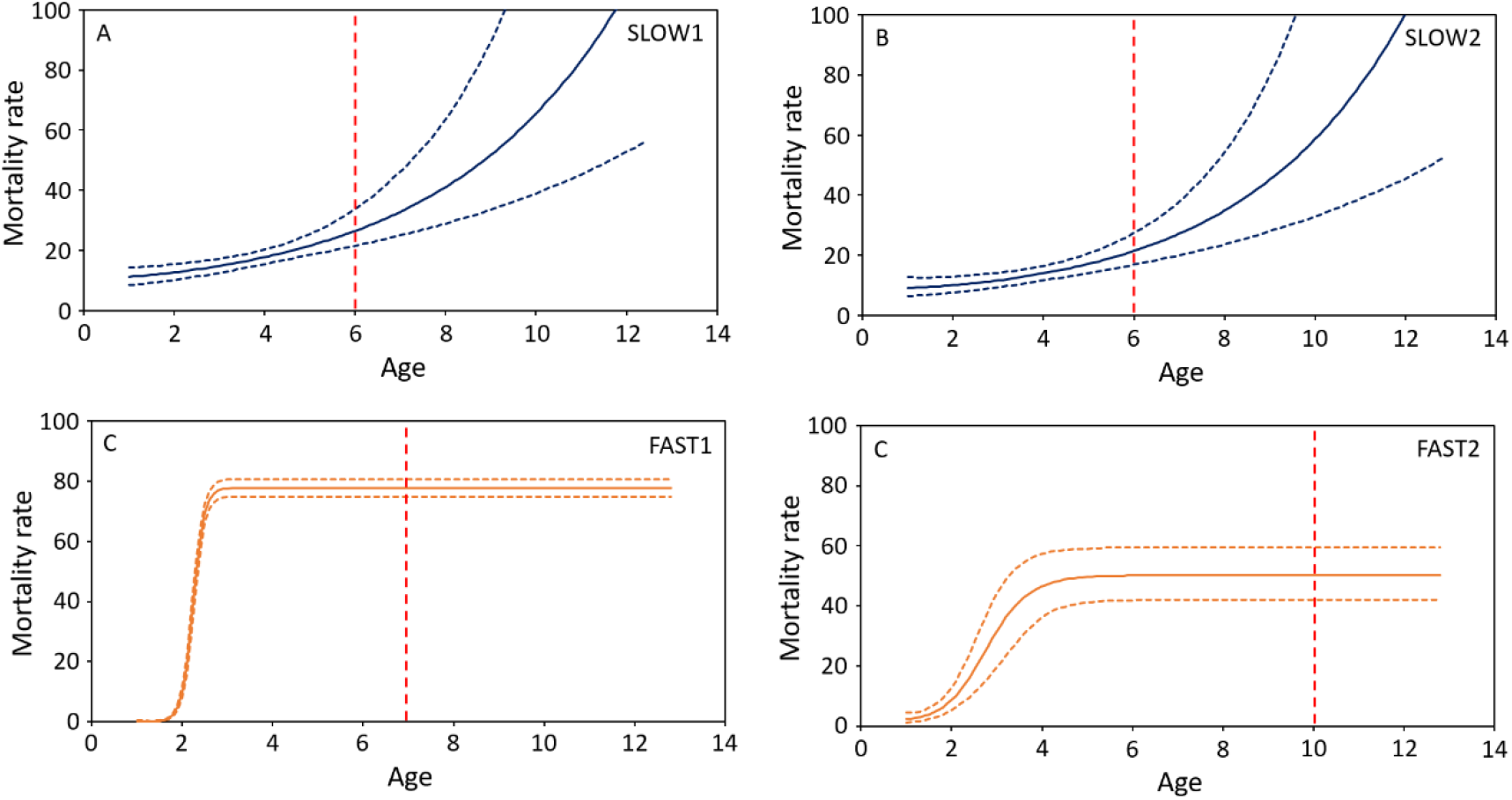
Mortality rate (i.e., proportion of individuals dying at a given age) in two fast (FAST1 and FAST2) and slow (SLOW1 and SLOW2) populations of *Bombina variegata*. The predictions on the left of the vertical dashed line correspond to observed ages while those on the right are model projections.

Age-dependent survival patterns were relatively similar between the populations with the same life history strategies but differed markedly between strategies (Fig. 3). In “slow” populations (Fig. 3A and 3B), the cumulative probability of surviving until a given age decreased slowly over a toad’s lifetime: it was 0.78 until age three, 0.43 until age six, 0.17 until age nine, and finally 0.00 until age 12 (Appendix 1, Table 2–3). In “fast” populations (Fig. 3C and 3D), cumulative survival probability decreased rapidly after two years: it was 0.58 until age three, 0.07 until age six, and 0.00 until age nine (Appendix 1, Table 4–5).

**Fig. 3.**
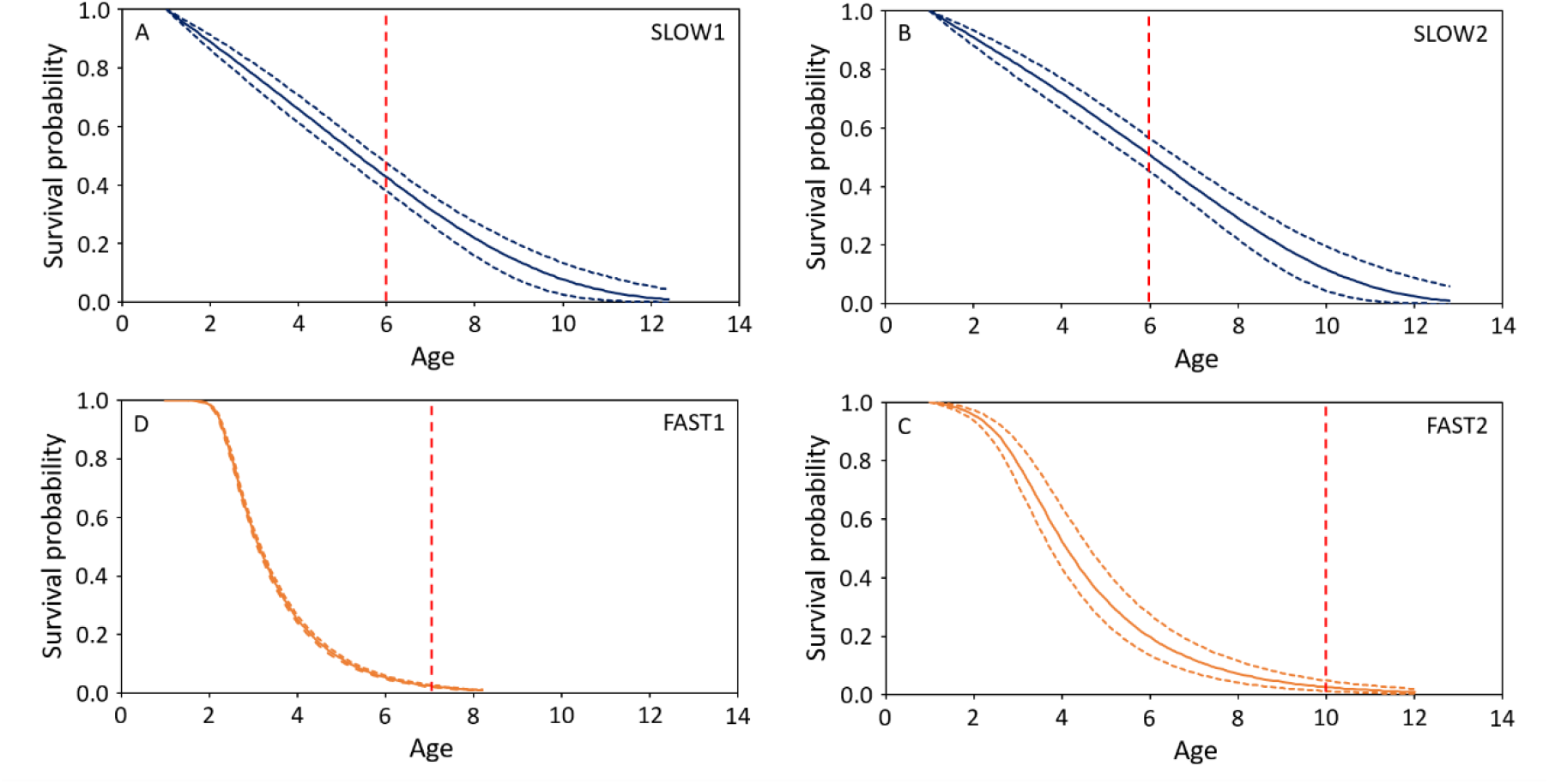
Survival probability until a given age in two fast (FAST1 and FAST2) and slow (SLOW1 and SLOW2) populations of *Bombina variegata*. The predictions on the left of the vertical dashed line correspond to observed ages while those on the right are model projections.

## Discussion

We provide the first clear evidence for actuarial senescence in a wild amphibian that is associated with an intraspecific slow-fast life history difference. The onset of senescence was earlier in individuals from “slow” populations and individuals aged at a faster rate than individuals in “fast” populations.

### Evidence for actuarial senescence in an amphibian

To date, senescence remains poorly studied in most ectothermic vertebrates due to the scarcity of demographic data with age records for these taxa (Conde et al. 2019). Amphibians are often absent from most publications investigating ageing across the tree of life (Ricklefs et al. 2010, Lemaître et al. 2015, Salguero-Gómez & Jones 2017). Lindström et al. (2010) reported an increase in age-specific mortality in a cave-dwelling salamander and Jones et al. (2014) provided evidence for senescence in a frog. Colchero et al. (2019) included anurans and salamanders in their comparative analyses and found a great diversity in patterns of senescence among the four species that were studied. Our analysis shows that there is also considerable variation at the intraspecific level. Our results suggest that there may be substantial variation in senescence within amphibians at both the interspecific and intraspecific level and that further exploration is worthwhile.

### The speed and shape of senescence depend on population-level pace-of-life

Jones and colleagues (2008) showed that interspecific variation in senescence can be explained by the position of the species along the slow-fast life-history gradient in mammals and birds (but see Jones et al. 2014 who showed that the relationship breaks down if the taxonomic range of the phylogenetic comparison is too broad [see also Stearns 1992 for a discussion of such broad life history comparisons]). The results presented here confirmed our prediction that the position on the fast-slow life history continuum affects actuarial senescence at the intraspecific level. Individuals from “fast” populations had an earlier onset of senescence (although our analysis did not allow a precise quantification of this onset) and aged at a faster rate than individuals from “slow” populations. In our study system, the factors (habitat type and latitude) causing the variation of life history strategies are potentially confounded: “fast” forest-dwelling populations were studied in northern France and “slow” river-dwelling populations in southern France. However, we assume that this confounding does not matter for the results that we present here – it could be problematic if one is interested in the processes that cause the life history differences, which was not the goal of our study. We also note that Cayuela et al. (2019) also found similar intraspecific variation of pace-of-life caused by the hydroperiod of breeding sites at the scale of a metapopulation. While it would be worthwhile to better understand the underlying causality, intraspecific variation along the slow-fast life-history gradient and the associated pattern of intraspecific variation in senescence are based on robust data and analyses.

Actuarial senescence likely evolves from an adjustment of resource allocation in response to environmentally-driven mortality (Kirkwood & Rose 1991; Stearns 2000, Baudisch & Vaupel 2012); In our study system, individuals from “fast” populations occurring in forest environment probably cope with environmentally-driven mortality risks associated with the unpredictability of breeding patches by allocating more resources to reproduction at the expense of somatic maintenance (see Cayuela et al. 2016a,b for an in-depth discussion). This might ultimately lead to a faster decline in individuals’ survival probabilities with age, as expected under the disposable soma theory of ageing (Kirkwood & Austad 2000). In contrast, environmentally-driven mortality is lower in river environment where the predictability of breeding resources is high. Accordingly, individuals may allocate fewer resources to reproduction – females produce a lower number of juveniles per year (Cayuela et al. 2016a) – and more to somatic maintenance, resulting in a later, slower senescence. A similar pattern was found in another ectothermic vertebrate, the western terrestrial garter snake (*Thamnophis elegans*; Robert & Bronikowski 2010). In this species, the ‘fast’ life-history type reaches a larger body size and has life-history traits that place it at the fast end of a fast-slow continuum (fast growth, early maturation, high reproductive output, shorter lifespan) relative to individuals of the small-bodied ‘slow’ life-history type. Studies highlighted that the individuals of the ‘slow’ life-history type, displaying a slower senescence, have DNA that damaged more readily but repaired more efficiently, and have more efficient mitochondria and more efficient cellular antioxidant defenses than individuals of the ‘fast’ ecotype (Robert & Bronikowski 2010).

### Conclusion

Our study and previous work suggest similarities in the macroevolutionary and microevolutionary processes shaping the evolution of ageing. We illustrate that the relationships between senescence patterns and the fast-slow life history continuum at the interspecific level can also be found at the intraspecific level. Our results suggest that the constraints imposed by trade-offs between fitness components appear to produce similar effects on ageing patterns at different levels of biological organization (i.e. intra- and interspecific levels). These results provide an important view of senescence in an ectothermic organism and contribute significantly to the broader study of senescence. First, intraspecific variation in senescence suggests that phylogeny should not strongly constrain the evolution of senescence as it has been conjectured (Shefferson et al. 2017). If evolution can lead to intraspecific variation in senescence (Stearns et al. 2000, Shefferson et al. 2017), then phylogenetic constraints would be a weak explanation for senescence patterns across the tree of life (Antonovics & van Tienderen 1991). Second, if there is intraspecific variation in senescence, this variation could explain the weakness of the phylogenetic signals in actuarial senescence patterns in several taxa (Salguero-Gómez & Jones 2017). It may well be that phylogenetic signals could become more apparent once intraspecific variation is taken into account. Such a combination of microevolutionary and macroevolutionary patterns of senescence would lead to a deeper understanding of the evolutionary biology of aging.

## Acknowledgments

We thank all the fiedworkers that have contributed to data collection. We also thank Erin Muths for her comments on the manuscript. This research was funded by the Lorraine DREAL, the Rhône-Alpes DREAL, the Agence de l’Eau Rhône-Alpes, the Agence de l’Eau Rhin-Meuse, the Office National des Forêts, the Conseil Régional de Lorraine, the Conseil Régional de Champagne-Ardenne, the Conseil Régional de Picardie, the Conseil Général de l’Aisne, the Conseil Général d’Ardèche, the Conseil Général d’Isère and the Communauté de Communes de l’Argonne Ardennaise (2C2A). Toad capture was authorized by the Préfecture de l’Ardèche (arrêté no. 2014–288-002) and the Préfecture de la Meuse (arrêté no. 2008–2150).

## APPENDIX 1: Additional information about *Bombina variegata* populations and capture-recapture data and analyses

**Fig. 1.**
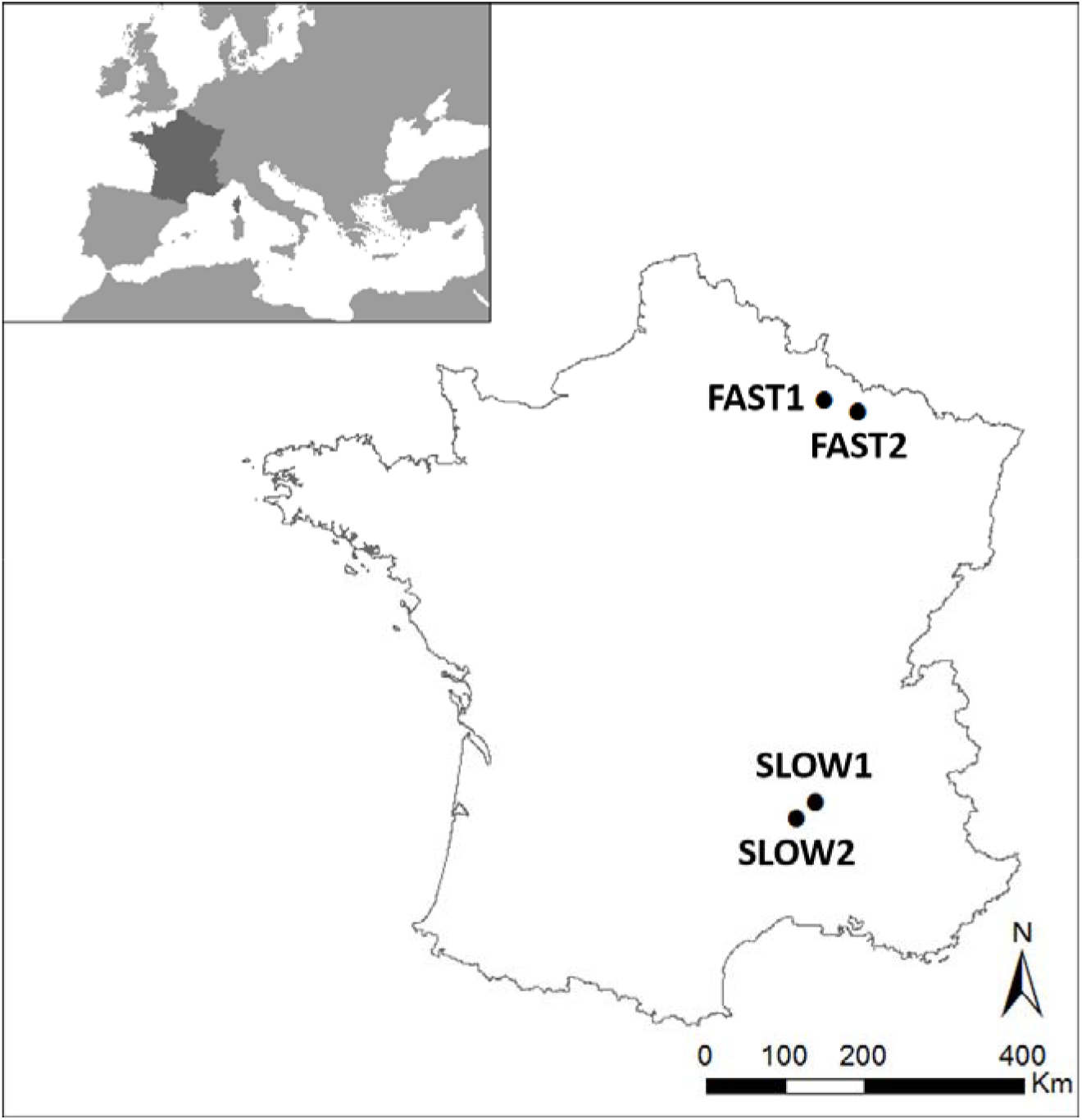
Localization of the two “slow” populations (SLOW1 and SLOW2) and the two “fast” populations (FAST1 and FAST2) in France.

**Fig.2.**
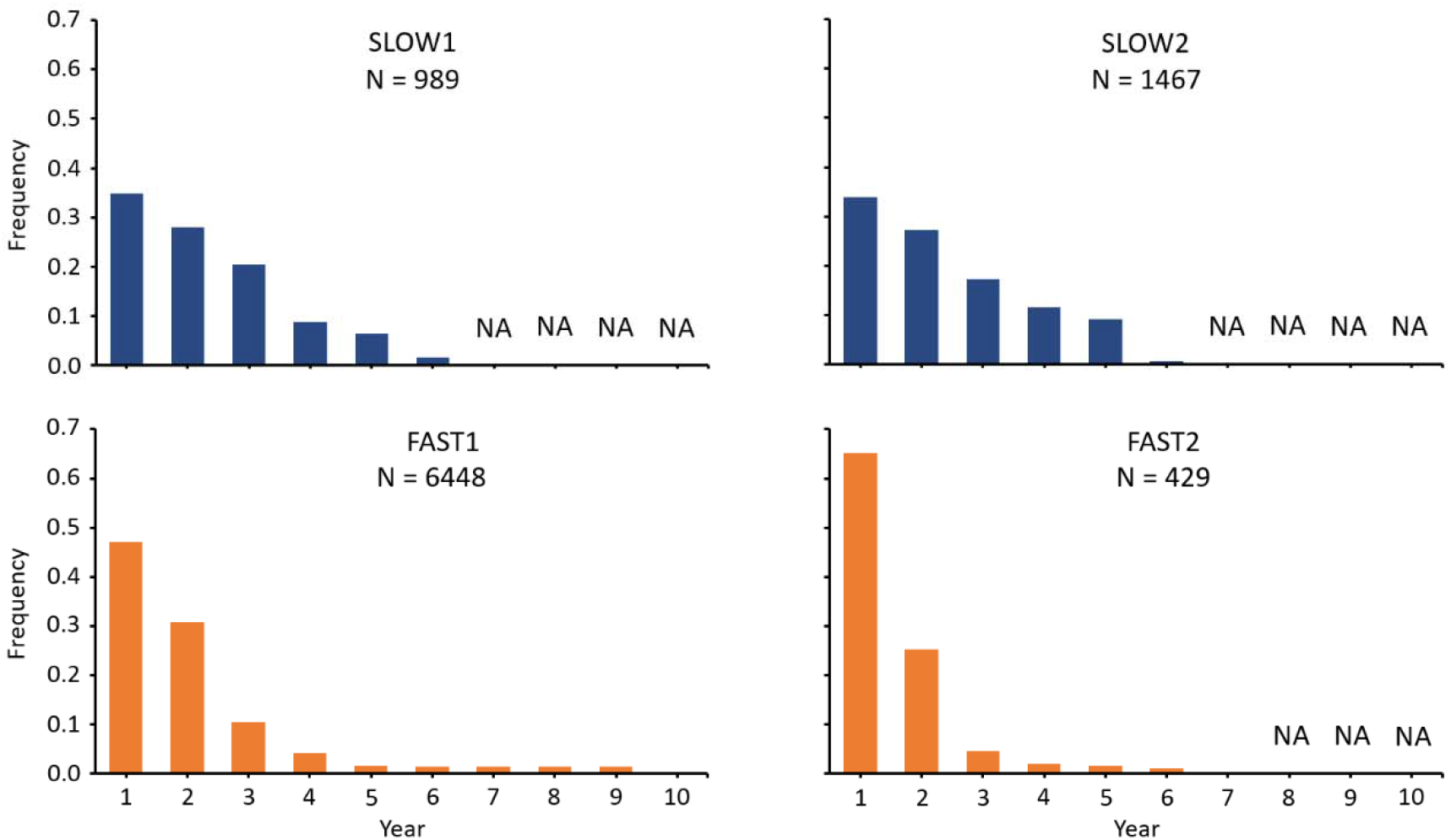
Age observations the two “slow” populations (SLOW1 and SLOW2) and the two “fast” populations (FAST1 and FAST2). NAs correspond to unavailable data for a given age class (due to survey length).

**Table 1.** Information about *B. variegata* populations (FAST1, FAST2, SLOW1 and SLOW2) and the capture-recapture data. Nb = number of individuals in the dataset, Year = number of years of survey, Sampling effort = number of capture sessions per year, Birth date = percentage of individuals for which the date of birth is known, Death date = percentage of individuals for which the date of death is known.

| Population | Life history | Nb | Year | Sampling effort | Birth date | Death date |
| --- | --- | --- | --- | --- | --- | --- |
| FAST1 | Fast | 580 | 9 | 3 to 12 | 51% | 0% |
| FAST2 | Fast | 9418 | 7 | 3 to 4 | 57% | 0% |
| SLOW1 | Slow | 1154 | 5 | 9 | 56% | 0% |
| SLOW2 | Slow | 768 | 5 | 9 | 53% | 0% |

**Table 2.** SLOW1: model outputs (Gompertz ‘bathtub’). Estimates, standard errors (StdErr) and 95% confidence intervals (95% CI) are provided. Survival: a0 = intercept, a1 = coefficient slope for age; mortality: b0 = intercept, b1 = coefficient slope for age, c = age-independent mortality parameter associated with the ‘bathtub’ function; recapture: pi = recapture probability for a given year.

| Coefficients | Estimate | StdErr | Lower95%CI | Upper95%CI |
| --- | --- | --- | --- | --- |
| a0 | -4.05673 | 0.686250 | -5.4091400 | -2.73486 |
| a1 | 0.36949 | 0.533950 | 0.0042349 | 1.97552 |
| c | 0.02554 | 0.020237 | 0.0009387 | 0.07434 |
| b0 | -2.80483 | 0.349414 | -3.6639895 | -2.26342 |
| b1 | 0.26081 | 0.057554 | 0.1514149 | 0.38224 |
| pi.2010 | 0.95596 | 0.008229 | 0.9385495 | 0.97099 |
| pi.2011 | 0.80983 | 0.016747 | 0.7746687 | 0.84140 |
| pi.2012 | 0.90937 | 0.011788 | 0.8848874 | 0.93129 |
| pi.2013 | 0.68960 | 0.017514 | 0.6550869 | 0.72326 |
| pi.2014 | 0.70550 | 0.021539 | 0.6634260 | 0.74761 |

**Table 3.** SLOW2: model outputs (Gompertz ‘bathtub’). Estimates, standard errors (StdErr) and 95% confidence intervals (95% CI) are provided. Survival: a0 = intercept, a1 = coefficient slope for age; mortality: b0 = intercept, b1 = coefficient slope for age, c = age-independent mortality parameter associated with the ‘bathtub’ function; recapture: pi = recapture probability for a given year.

| Coefficients | Estimate | StdErr | Lower95%CI | Upper95%CI |
| --- | --- | --- | --- | --- |
| a0 | -3.99578 | 0.64023 | -5.310219 | -2.83556 |
| a1 | 0.49436 | 0.61439 | 0.005362 | 2.27773 |
| c | 0.02726 | 0.02039 | 0.001082 | 0.07551 |
| b0 | -3.22119 | 0.47976 | -4.438314 | -2.54104 |
| b1 | 0.29107 | 0.07283 | 0.168434 | 0.46047 |
| pi.2010 | 0.93446 | 0.01281 | 0.907488 | 0.95797 |
| pi.2011 | 0.83983 | 0.01807 | 0.803634 | 0.87304 |
| pi.2012 | 0.78004 | 0.02296 | 0.732969 | 0.82302 |
| pi.2013 | 0.81596 | 0.01989 | 0.775446 | 0.85390 |
| pi.2014 | 0.83182 | 0.01888 | 0.793399 | 0.86712 |

**Table 4.** FAST1: model outputs (logistic ‘bathtub’). Estimates, standard errors (StdErr) and 95% confidence intervals (95% CI) are provided. Survival: a0 = intercept, a1 = coefficient slope for age; mortality: b0 = intercept, b1 = coefficient slope for age, c = age-independent mortality parameter associated with the ‘bathtub’ function; recapture: pi = recapture probability for a given year.

| Coefficient | Estimate | StdErr | Lower95%CI | Upper95%CI |
| --- | --- | --- | --- | --- |
| a0 | -4.646298 | 0.556074 | -5.823e+00 | -3.62562 |
| a1 | 1.246570 | 0.757036 | 9.695e-02 | 2.94712 |
| c | 0.003002 | 0.003066 | 6.853e-05 | 0.01129 |
| b0 | -4.558240 | 0.361952 | -5.295e+00 | -3.89086 |
| b1 | 2.175048 | 0.340004 | 1.569e+00 | 2.87970 |
| b2 | 4.374321 | 0.657421 | 3.113e+00 | 5.69759 |
| pi.2000 | 0.969228 | 0.019928 | 9.217e-01 | 0.99632 |
| pi.2001 | 0.762933 | 0.036631 | 6.880e-01 | 0.83264 |
| pi.2002 | 0.606212 | 0.049487 | 5.089e-01 | 0.70316 |
| pi.2003 | 0.311041 | 0.049952 | 2.215e-01 | 0.41663 |
| pi.2004 | 0.167734 | 0.039902 | 9.900e-02 | 0.25432 |
| pi.2005 | 0.222645 | 0.038082 | 1.534e-01 | 0.29990 |
| pi.2006 | 0.568990 | 0.036065 | 4.959e-01 | 0.63895 |
| pi.2007 | 0.575538 | 0.029580 | 5.153e-01 | 0.63211 |
| pi.2008 | 0.723976 | 0.035850 | 6.551e-01 | 0.79512 |

**Table 5.** FAST2: model outputs (logistic ‘bathtub’). Estimates, standard errors (StdErr) and 95% confidence intervals (95% CI) are provided. Survival: a0 = intercept, a1 = coefficient slope for age; mortality: b0 = intercept, b1 = coefficient slope for age, c = age-independent mortality parameter associated with the ‘bathtub’ function; recapture: pi = recapture probability for a given year.

| Coefficient | Estimate | StdErr | Lower95%CI | Upper95%CI |
| --- | --- | --- | --- | --- |
| a0 | -6.514e+00 | 0.4849332 | -7.519e+00 | -5.5936923 |
| a1 | 1.732e+00 | 0.7944760 | 2.886e-01 | 3.3760323 |
| c | 1.961e-04 | 0.0001904 | 5.350e-06 | 0.0007034 |
| b0 | -1.026e+01 | 0.3220379 | -1.091e+01 | -9.6254164 |
| b1 | 8.076e+00 | 0.2864652 | 7.513e+00 | 8.6570983 |
| b2 | 1.040e+01 | 0.4136846 | 9.602e+00 | 11.2350077 |
| pi.2008 | 9.095e-01 | 0.0127453 | 8.844e-01 | 0.9341675 |
| pi.2009 | 5.379e-01 | 0.0094210 | 5.196e-01 | 0.5566322 |
| pi.2010 | 5.455e-01 | 0.0097414 | 5.267e-01 | 0.5646473 |
| pi.2011 | 1.629e-01 | 0.0073179 | 1.489e-01 | 0.1774715 |
| pi.2012 | 5.793e-01 | 0.0099371 | 5.597e-01 | 0.5989854 |
| pi.2013 | 5.938e-01 | 0.0081348 | 5.780e-01 | 0.6102405 |
| pi.2014 | 6.809e-01 | 0.0078857 | 6.653e-01 | 0.6961301 |

